# Soleus H-reflex size versus stimulation rate in the presence of background muscle activity: A methodological study

**DOI:** 10.1101/2025.03.17.643784

**Authors:** Jodi A. Brangaccio, Disha Gupta, Helia Mojtabavi, Russell L. Hardesty, N. Jeremy Hill, Jonathan S. Carp, Darren E. Gemoets, Theresa M. Vaughan, James J. S. Norton, Monica A. Perez, Jonathan R. Wolpaw

## Abstract

**Introduction:** Hoffmann reflex (HR) operant conditioning has emerged as an important intervention in neurorehabilitation. During conditioning, the HR is elicited at low rates (∼0.2 Hz) to avoid the initial reduction in HR size that can occur over repeated stimulation, i.e., rate-dependent depression (RDD), thereby maintaining reflex size. This study investigated the impact of higher stimulation rates on HR size, where a stable, low-level, background electromyographic (EMG) signal is maintained over 225 conditioning trials in each of 30 sessions. A higher rate could shorten session length and/or number.

**Methods:** Fifteen healthy participants maintained low background soleus EMG (5-18 µV, ∼1-3% of the maximum stimulation evoked direct muscle (M-wave) EMG response (M_max_) while standing. Soleus HR and M-wave recruitment curves were obtained at rates of 0.2, 1, and 2 Hz, from which M_max_ and H_max_ were calculated. Seventy-five HR trials (HRT) were collected for each stimulation rate at a target M-wave size (∼10-20% of M_max_).

**Results:** There was no evidence of RDD at higher stimulation rates. In addition, the mean HR over trials was reliable across participants and rates. The Intraclass Correlation Coefficient (ICC) was 0.965 (95%CI:0.915, 0.987).

**Discussion:** This study shows that H-reflex conditioning might be performed at rates up to 2 Hz with no RDD and with consistent HR values. A faster rate could increase the number of conditioning trials per session, reduce session duration, and/or reduce the number of sessions. It could thereby accelerate the conditioning process and make the process less demanding for participants.

**Support:** NIH Grant P41 EB018783 (Wolpaw), NYS Spinal Cord Injury Research Board C37714GG (Gupta) and C38338GG (Wolpaw), VA SPiRE NCT05880251(Brangaccio), Stratton Veterans Affairs Medical Center.

## Introduction

Operant conditioning of spinal reflexes is a targeted noninvasive therapy for restoring motor skills impaired by neurological injury or disease (e.g., stroke, spinal cord injury) (Kim et al., 2023; Thompson et al., 2022, 2019b, 2013). During H-reflex operant conditioning (HROC), participants learn to change a spinal reflex behavior (Kim et al., 2023; Makihara et al., 2014; Thompson et al., 2022, 2019b, 2013, 2009). This targeted beneficial plasticity, when paired with skill-specific practice, can lead to widespread multi-site plasticity in the central nervous system (CNS), thereby enabling overall improvements in motor control (e.g., walking, balance, mobility) in people with functional deficits due to neurological disorders (Kim et al., 2023; Thompson et al., 2022, 2019a, 2013).

The ability of HROC to promote targeted beneficial plasticity at key sites make it a meaningful addition to traditional skill-specific practice (Wolpaw et al 2023). However, the several months required for effective HROC. Current protocols require up to 36 one-hour sessions three times per week over 12 weeks; each session comprises 225 trials or a total of 8100 stimuli. Adding more trials by lengthening session duration is impractical from a clinical standpoint. Thus, we are investigating increased stimulation rate as a way to increase the dose of HROC and reduce the time it takes to realize meaningful functional change.

At stimulation rates >0.2 Hz, H-reflex (HR) size undergoes rate-dependent depression (RDD) (Lloyd and Weber, 1957; Ishikawa, et al., 1966). RDD can be assessed in healthy humans and animals and reflects the action of spinal inhibitory systems (Capek and Esplin 1977, Chang, et al 2020, Hultborn et al, 1996, Lee-Kubli, et al 2014). It is believed to be important in regulation of the stretch reflex during dynamic conditions, e.g. walking and standing (Capaday, et al., 1986).

Our current soleus muscle (SOL) HROC protocol (Hill et al., 2022) and previous studies (Makihara et al., 2014; Thompson et al., 2013, 2009) applied stimulation at ≤0.2 Hz to minimize inducing RDD. RDD is prominent when background EMG is absent; it is less evident when background EMG activity is present (Burke et al., 1989; Clair et al., 2011; Crone and Nielsen, 1989; McNulty et al., 2008; Rothwell et al., 1986; Stein et al., 2007). In HROC conditioning, immediate feedback of HR size guides the patient in learning to change HR size in the beneficial direction. Therefore, an HR that is both large enough and stable enough to provide reliable and accurate feedback is essential for effective HROC. The goal of this study was to determine whether our HROC protocol can be performed at rates higher than 0.2 Hz without causing significant RDD that might impair this essential feedback.

We hypothesized that the low level of SOL background EMG that our participants are required to maintain during the HROC protocol would be sufficient to suppress RDD at stimulation rates higher than 0.2 Hz. An ability to increase the number of trials per session without inducing RDD may allow HROC to produce the same or even greater beneficial effects with fewer conditioning sessions, and lead to a more nuanced understanding of HROC dosing.

## Methods

### Participants

Fifteen neurologically intact participants (mean age 51±19 SD years, 7 females) gave informed consent prior to participating in the study. The study was performed at the Albany Stratton VA Medical Center, Albany, NY and approved by the Albany Stratton IRB. Inclusion criteria included: (1) ≥18 years old; (2) no known history of neurological disease; (3) no significant cardiac history; and (4) measurable SOL HR during standing. Prior to beginning the study, participants were randomly assigned to one of three specific sequences in the order of the three stimulation rates (Suppl Table S1 for participant profiles and sequence.)

### EMG Recordings and Electrical Stimulation

At the beginning of each session, both legs were cleaned with alcohol and gently abraded prior to the application of EMG recording and stimulating electrodes. For recording EMG, surface Ag-AgCl electrodes were placed longitudinally in bipolar fashion over soleus and tibialis anterior (TA) muscles bilaterally approximately 3 cm apart center-to-center (Makihara et al., 2014; Thompson et al., 2013, 2009). EMG was amplified (total gain=500) and bandpass filtered (10–1000 Hz) (AMT-8 EMG amplifier, Bortec Biomedical Ltd., Calgary, Alberta, Canada), and then digitized (3200 Hz) and stored using our evoked potential operant conditioning software (EPOCS) (Hill et al., 2022).

The SOL HR was elicited by transcutaneous electrical stimulation (DS8R stimulator, Digitimer Ltd, Welwyn Garden City, UK) using a 1-ms square biphasic waveform applied through surface Ag-AgCl electrodes (2.2 × 2.2 cm for the cathode and 2.2 × 3.5 cm for the anode and ground; Vermed) with the cathode over the popliteal fossa crease and the anode 5-7 cm rostral to the cathode. Electrode position was then optimized to minimize the stimulation current required to elicit a threshold HR, while maximizing HR size and participant comfort. The ground electrode was on the patella.

We quantified background EMG level (BG), direct muscle response (M-wave) size, and HR size as mean rectified amplitude within a preselected time window specific to each participant. Generally, these windows (relative to the time of stimulus delivery) were -52 to -2 ms (pre-stim) for BG; 7 to 20 ms for the M-wave; and 35 to 50 ms for the HR (Hill et al., 2022; Thompson et al., 2009).

### Experimental Design

The aim of this study was to determine whether our current HROC procedures (Hill et al., 2022) could be successfully implemented at rates higher than our standard 0.2 Hz without significant RDD. To do this, we performed two stimulation procedures that are key components of our HROC protocol at rates of 0.2, 1.0, and 2.0 Hz. The first was the Recruitment Curve (RC) procedure, in which we measure HR size and M-wave size over a wide range of stimulation intensities from their thresholds to their maxima. Using this data we generated recruitment curves to assess the maximum M-wave and HR (M_max_ and H_max_, respectively), and determined an M-wave target value (Thompson et al., 2013, 2009). The second procedure was the H-reflex trials (HRT) procedure, in which we measured M-wave and HR sizes over a block of 75 H-reflex trials (HRTs) with a fixed M-wave target (typically just above the threshold M-wave value and below the M-wave value at H_max_). The data from the HRT procedure was then used to evaluate the effect of stimulation rate on RDD.

All measurements were made in one 120-min session. A session included two runs at each of the three frequencies, for a total of six runs. Each run comprised the RC and HRT procedures. To randomize the six different orderings of three stimulation frequencies for each participant, the stimulation frequencies were counterbalanced in a pseudo-randomized block of six, e.g., if the first run was 0.2, 1.0, and 2.0 Hz, the second run was 2.0, 1.0, and 0.2 Hz. (See Table 1 for an example, and Table S1 for the ordering assigned to each participant.) The interstimulus interval varied randomly within 20% of its mean to prevent the participant from anticipating stimulus delivery time. The same people conducted all study phases to avoid inter-operator variability.

**Table 1.**
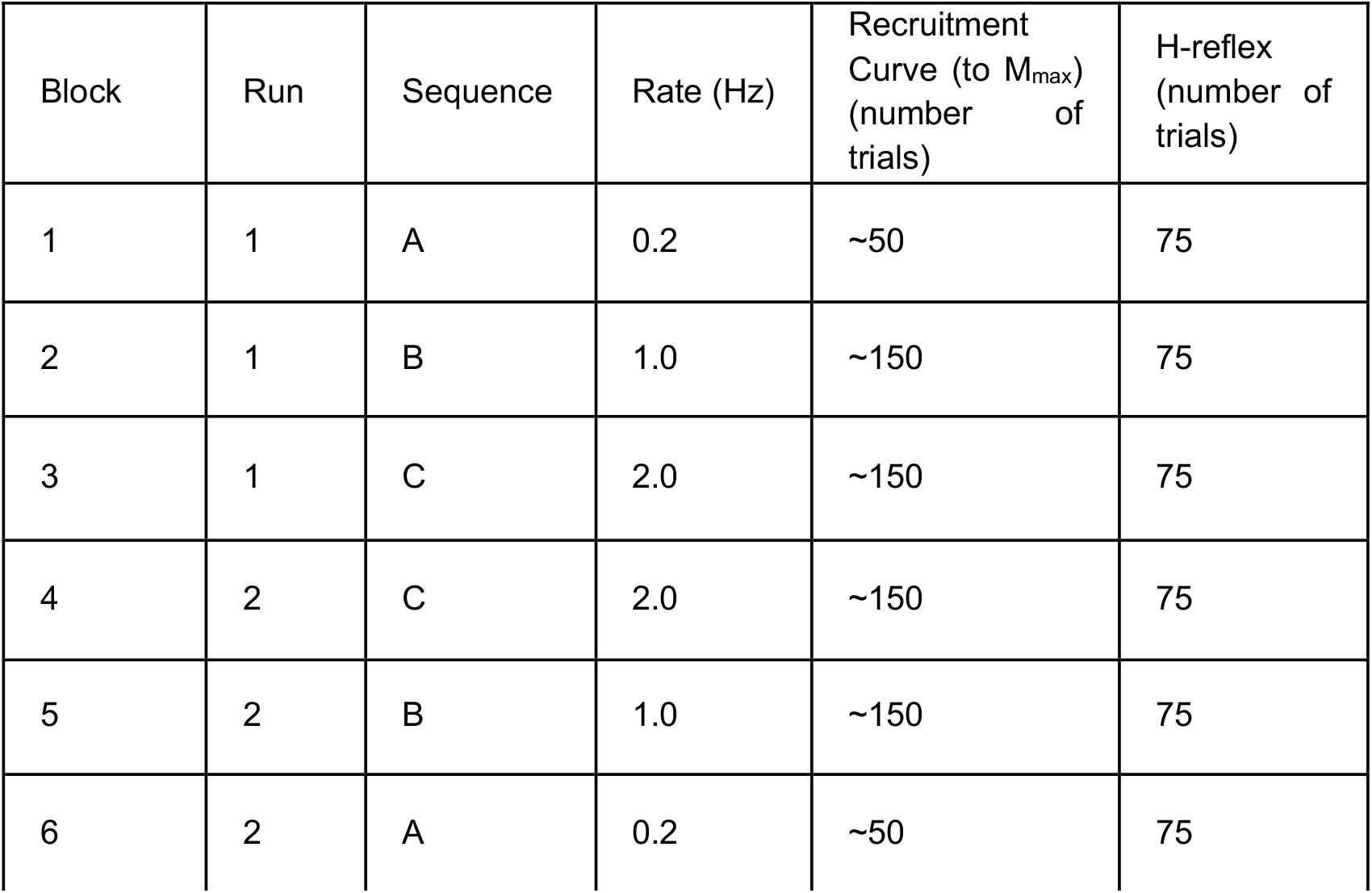
An example of the experimental design in one participant. Six runs of the recruitment curve (RC) procedure and the H-reflex trial (HRT) procedure were performed at a pseudo-randomized block of three different rates (*A, B*, and *C*) where the values of *A, B*, and *C* reflect randomized sequences of 0.2 Hz, 1 Hz, and 2 Hz. Run 1 was performed using one of three sequences: ABC; BCA; CAB. Run 2 was performed after a brief sitting break by repeating the RC and HRT procedures in the reverse order of rates used in Run 1 (e.g., CBA for ABC; see Suppl Table S1 for all sequences). The interstimulus interval used was randomly varied within 20% of its mean to minimize the chance of the participant anticipating the exact stimulus delivery time.

| Block | Run | Sequence | Rate (Hz) | Recruitment Curve (to $M_{\max}$) (number of trials) | H-reflex (number of trials) |
| --- | --- | --- | --- | --- | --- |
| 1 | 1 | A | 0.2 | ~50 | 75 |
| 2 | 1 | B | 1.0 | ~150 | 75 |
| 3 | 1 | C | 2.0 | ~150 | 75 |
| 4 | 2 | C | 2.0 | ~150 | 75 |
| 5 | 2 | B | 1.0 | ~150 | 75 |
| 6 | 2 | A | 0.2 | ~50 | 75 |

**Table 2.**
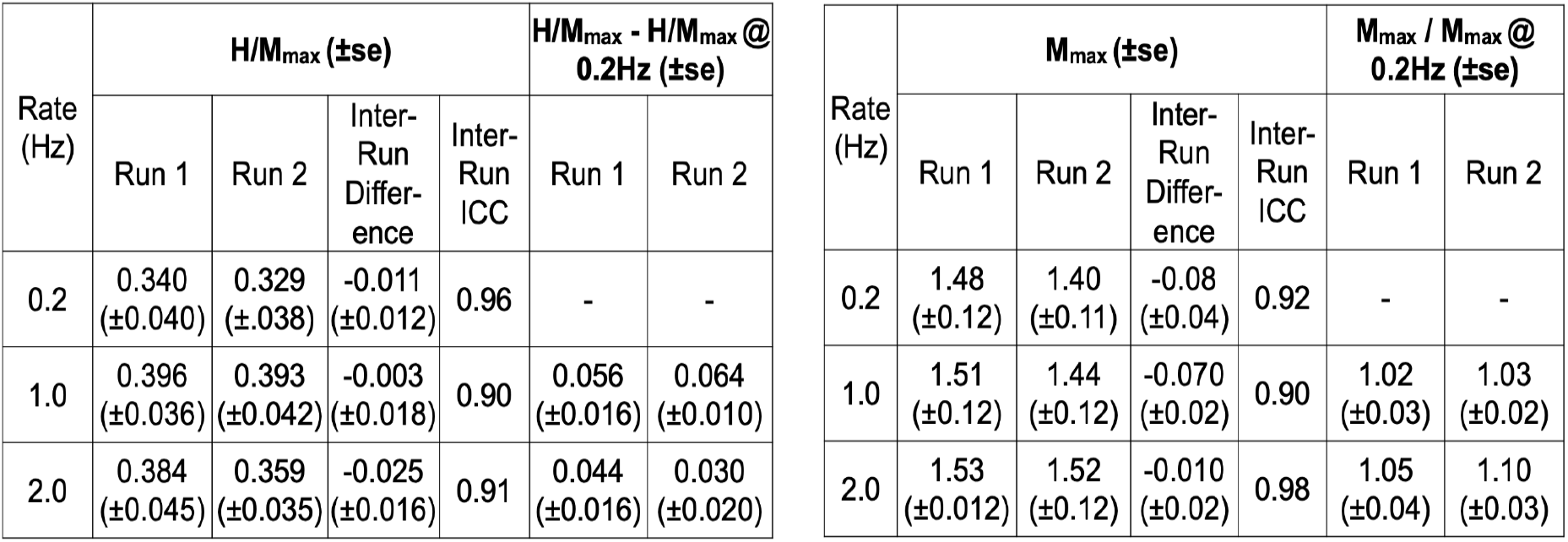
Comparison of the RC data across rates and runs. The table displays the inter-run differences and ICCs within M_max_ and H_max_ (normalized to M_max_) across rates and runs. There were no significant differences found in M_max_ and H_max_ between Runs 1 and 2 or between stimulation rates of 1 or 2 Hz and 0.2 Hz (p>0.3 for all inter-run differences).

### Recruitment Curve Protocol

Measurements were made with the participant maintaining a natural standing posture and looking forward at a large monitor for visual feedback. If needed for balance, the participant was given the minimal upper extremity support of a walker. The EMG responses to tibial nerve stimulation were recorded at the appropriate rate while the participant maintained a modest level of soleus EMG activity, equal to natural static standing. The mean rectified BG was in the range of 5-18 µV (∼1-3% of their M_max_). Antagonist TA activity was also recorded and maintained (∼0-10 µV). Stimulus intensity was increased from the HR threshold to the maximum H-wave (H_max_) and then to an intensity just above that needed to elicit the maximum M-wave (M_max_) (Kido et al., 2004; Thompson et al., 2009; Zehr and Stein, 1999).

The recruitment curves were performed as follows For the 1 and 2-Hz rates, stimulation intensity was continuously increased by 0.1-0.3 mA throughout the recruitment curve until the HR and M-wave maxima were determined. This required up to 150 trials (∼1.5-3.0 min). To determine HR and M-wave maxima we averaged the four adjacent trials at the peak. For the 0.2-Hz rate, we performed the RC trials with our standard methodology of stimulating every four trials at the same intensity and incrementing by ∼1.5 mA until the HR and M-wave maxima were reached, which usually occurred in about 50 trials (∼4-5 min). To determine HR and M-wave maxima we averaged the four trials at the maximum.

### H-Reflex Trial (HRT) Procedure

For the 75 HRT, a target M-wave size was selected based on that rate’s corresponding recruitment curve using three criteria such that the M-wave target was: (1) above M-wave threshold; (2) on the ascending slope of the HR RC where the M-wave and HR were both increasing; and (3) below H_max_, generally this is ∼10-20% of M_max_ (Crone, et al 1986). The M-wave target is a key component of all of our human and animal HROC studies; it helps to ensure that, throughout the multiple sessions of the HROC protocol, the same group of motoneurons (and by extension the same group of primary afferent fibers) is excited by the stimulus that evokes the HR.

During the HRT procedure, the stimulus current was manually adjusted as needed to maintain M-wave size within 20% of the target M-wave value. BG, M-wave size, and HR size were quantified as the mean rectified EMG amplitude in their time windows. These values were divided by M_max_ from the corresponding recruitment curve to normalize between participants, and thereby control for inter-person variability.

### Data and Statistical Analysis

#### Recruitment Curve Procedure

To assess differences and consistency in recruitment curve features (M_max_, H_max_/M_max_, and BG) within and between participants across runs, and across rates, we used linear mixed-effects models and ICCs (two-way ANOVA with main effects of run and rate and their interaction, and a within-subject repeated measures design)

#### H-reflex Trial Procedure

We expected RDD to manifest as a nonlinear decrease in mean rectified HR over the first few trials across participants. Thus, to test for RDD, we fit mixed-effects regression models to the first 20 trials assuming a nonlinear trend. For fixed effects, we included a three-way interaction between run/rate/trial, all two-way interactions, and main effects. Random effects were included allowing subject-specific intercepts and nonlinear trends. Likelihood-ratio tests were used to determine the significance of any fixed effects.

To estimate participant-specific level of effect on the magnitude of RDD, we fit analogous nonlinear regression models to each participant and summarized the model parameter estimates.

The aforementioned mixed-effects models tested the existence of RDD across all participants (i.e., in the population). We also quantified possible within-participant RDD over all 75 trials by fitting linear regression models separately for each participant within each run/rate combination. The slope values were summarized by boxplots expressed as percent change in mean rectified HR over the 75 trials (Supplemental Fig S6).

To assess reliability in the HRT across participants, we averaged over trials within subject/run/rate to remove the trial effect, then refit the mixed-effects models previously described. After determining no presence of run-by-rate interaction, we used ICCs for assessing absolute agreement for run and rate separately, assuming random rows and columns (i.e., random subjects and random treatment effects, usually denoted ICC2). (Matthias Gamer, et al., 2005; R Core Team, 2024)

The mixed-effects models employed the R package *lme4*, and the ICC values were computed using the package *irr* (Bates et al., 2015; Matthias Gamer, et al., 2005).

## Results

This study sought to determine whether the HR measurements performed during our HROC protocol could be elicited at rates as fast as 2 Hz without significant RDD or other effects on HR size. To verify our results, we first ran linear mixed-effects models and ICCs on the recruitment curve measurements for each participant, run, and rate. Once consistency and reliability were confirmed in the RC, we began our analyses for the presence of RDD within the HRT. Given that RDD is typically most pronounced in the initial trials of continuous trains of stimulation,we analyzed the first 20 trials of the HRT looking for a decreasing linear trend over stimulation repetition, effect of run or effect of rate, including all possible interactions. There was no linear trend over stimulation repetition, effect of run or effect of rate; thus, there was no evidence of RDD. Next, we analyzed all 75 HRT for repeatability, consistency, differences and trends within and across rates, runs, trials, and participants. Mixed-effects regression models, ICCs, and box plots were used and are explained in detail below.

### Recruitment Curve Results

Prior to investigating the presence of RDD in the HRT, the M-wave and HR recruitment curves were obtained, allowing us to confirm consistency and reliability in the curves across runs, rates, and participants. The recruitment curves were all similar in shape and latency. Figure 1 shows a typical example from one participant: the M-wave and HR recruitment curves are similar across rates and runs. Hypothesis testing showed no significant differences in M_max_ and H_max_/M_max_ between runs or rates (p>0.3), and all the ICCs were ≥ 0.90 between runs and rates (see Table 3), confirming the similarity of the RCs over all participants and validating the use of the M-wave RC to determine the M-wave target for the HRT.

**Figure 1.**
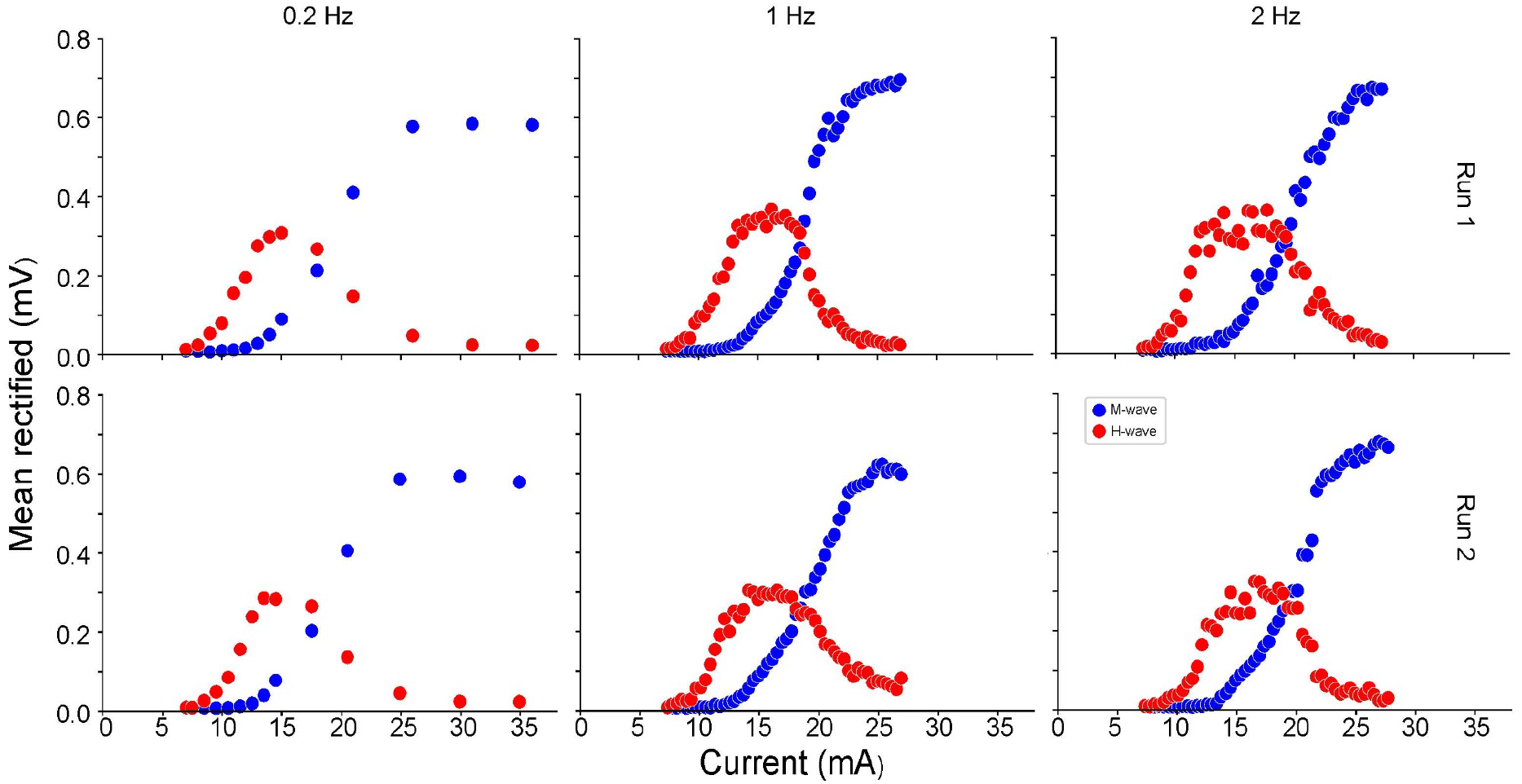
Example of the M-wave (blue) and HR (red) recruitment curves for each rate and run from one subject.

### HRT Results (H-Reflex Measurements)

#### Assessment of nonlinear trends over first 20 trials

The HRT Procedure was always performed after a period of rest (a seated break), after performing the RC Procedure for that rate. It was hypothesized that any RDD would likely occur within these first 20 HRT. We expected the expression of RDD to be nonlinear, likely an exponential decay to a steady-state value. Plots of the raw data (Supplemental Figure S1) offered no evidence of a decrease over the first 20 trials across all participants; this is supported by mixed-effects models which also found no significant nonlinear decrease (p=0.31, see “Mixed-effect model details” in Supplemental Materials for further information).

The mixed-effects models assessed statistical significance for any presence of RDD across the participants. Even though none was found, we also estimated effects of RDD at the participant-level. Figure 2 presents box plots summarizing the exponential rate parameter (i.e., a in the nonlinear model *exp(ax)*, where *a* is the exponential decay constant and *x* is the trial number) for the 15 participants. Note that an estimate of a<0 would be consistent with the expression of RDD (i.e., HR decreases exponentially over trials); a value of *a*=0 would yield a constant value for exp(*a*x) (i.e., no RDD). The boxplots are comparable across runs and rates, both in location (with median estimates (solid line in middle of the box) located near 0), and in distribution (box represents the interquartile range).

**Figure 2.**
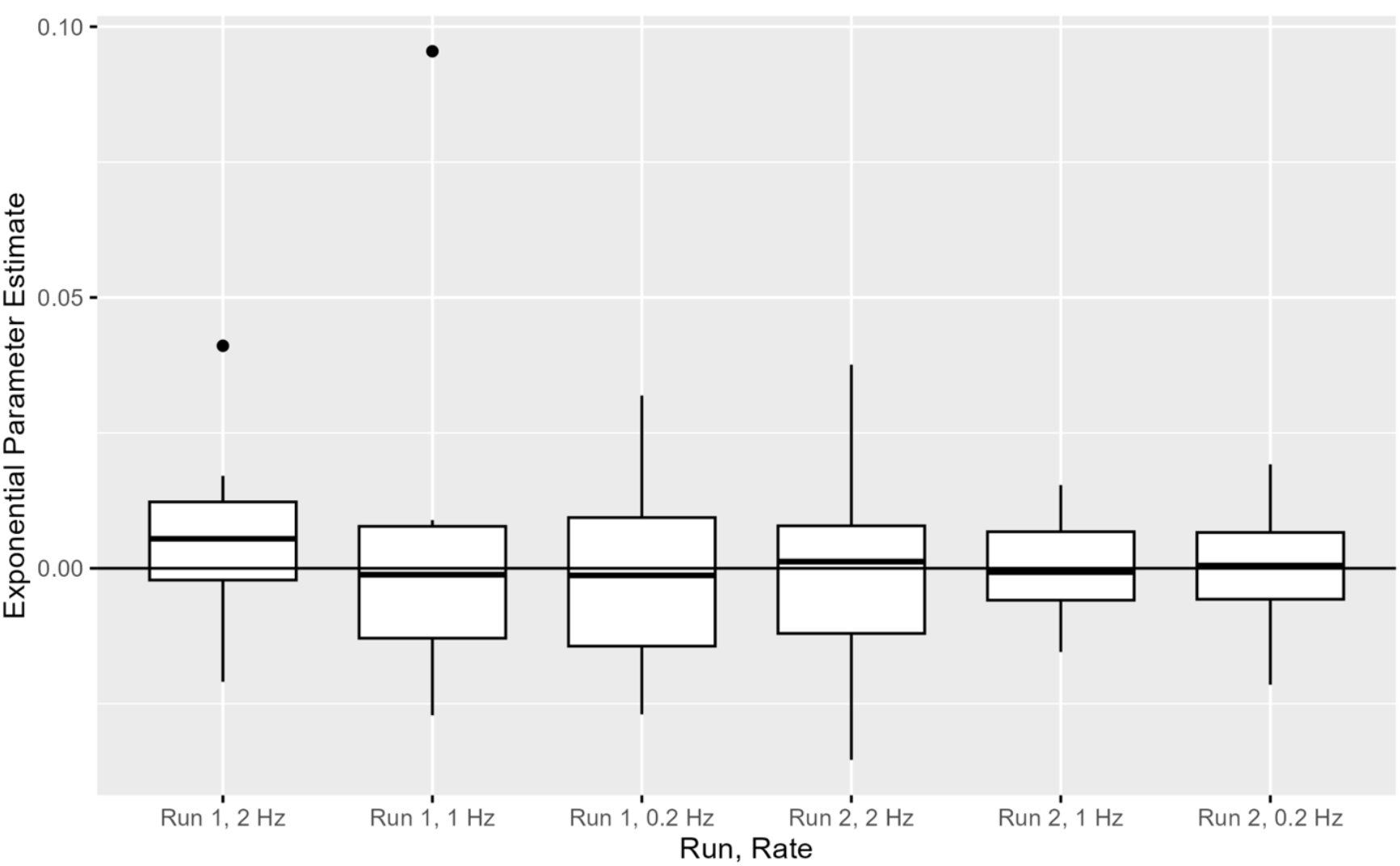
Participant-level estimates of the parameter a in the exponential model f(x)=b exp(ax) fit to each person in the mixed-effects model, where x is the trial number and a is the exponential parameter estimate. Boxplots summarize the estimates for the 15 participants. An estimate of a<0 would suggest RDD (i.e., HR decreases exponentially over trials). The boxplots are comparable across runs and rates, with median estimates (solid line inside of the box) located near *a*=0. Note that exp(0)=1, a constant value.

#### Testing for differences over all 75 Trials

After looking at the first 20 trials, to determine the presence of RDD, we then used hypothesis testing and analyzed the 75 H-reflex trials across rates and runs. For BG, we found no significant effect of rate, run or trial (see Supplemental Table S2 in the for the full analysis, and Supplemental Figure S4 for a plot of the raw data) (i.e., BG is essentially constant for all participants across trial/run/rate and is not inadvertently being used to change or affect the HR value). Supplemental Figure S5 (right panel) shows BG rates averaged over run and trial for each participant.

For HR values, there were no significant differences between runs or across trials (assuming linear trend over trials). However, there were modest but significant differences in mean HR between the rates 1 Hz and 0.2 Hz (p=0.002) and 2 Hz and 0.2 Hz (p=0.006). The relative increases in HR values at 0.2 Hz (mean value 0.259) compared to 1 Hz (mean value 0.231) and 2 Hz (mean value 0.233) are 10.0% and 10.8%, respectively. Figure S5 (left panel) shows normalized H values averaged over trial and run. The full statistical analysis is in the Supplemental, and plots of the raw data in Figure S2. This percent decrease in overall HR was consistent with findings of previous studies (Stein, et al 2007)

For M-wave values, one significant difference (p=0.02) was found between 1 Hz (mean value 0.121) and 0.2 Hz (mean value 0.106); this was a 14.2% relative increase in M-wave values in 1 Hz vs 0.2 Hz. A full description of these statistical analyses is in the Supplemental “Mixed-model effect details,” and plots of the raw data are in Supplemental Figure S4.

#### Assessment of consistency and agreement over all 75 Trials

After establishing minimal change in HR, M-wave, or BG over the 75 HRTs (Figure 2), we then calculated averaged values of HR, M-wave, and BG over the 75 HRTs. Confirming no evidence of run-by-rate interaction effects using mixed-effects models on these averaged data, we then computed ICCs by run and rate separately (Table 3). All but one of the resulting ICC values are >0.95, supporting high consistency. The ICC for the effect of rate for BG EMG is slightly lower but still nearly 0.9. These values further support the consistency in our HR values and lack of influence or control from muscle activation or procedural instability.

**Table 3.** Contains ICC values (95% CI) using the mean trial value within each subject/run/condition. We are confident in reporting ICCs averaged over trials because there are only minimal trends in trials across participants and treatment conditions. All three measures, BG, M-wave, and HR, have very high ICC values (an ICC value of 1 would indicate perfect agreement), with the lowest estimate of 0.889 for BG/rate. For BG rate, the value of 0.889 still indicates very good reliability (Koo and Li, 2016).

| Variable | Treatment | ICC (95% CI) |
| --- | --- | --- |
| HR size (normalized to $M_{\max}$) | Run | 0.987 (0.964,0.996) |
|  | Rate | 0.965 (0.915, 0.987) |
| M-wave size (normalized to $M_{\max}$) | Run | 0.985 (0.957, 0.995) |
|  | Rate | 0.978 (0.949, 0.992) |
| BG (normalized to $M_{\max}$) | Run | 0.966 (0.864, 0.990) |
|  | Rate | 0.889 (0.762,0.957) |

## Discussion

Up to now, HROC studies have been performed with a stimulation rate of 0.2 Hz or slower to avoid eliciting RDD (Makihara et al., 2014; Thompson et al., 2013, 2009). However, RDD can be suppressed by low levels of muscle activity associated with active movement or standing (Burke et al., 1989; Clair et al., 2011; Crone and Nielsen, 1989; McNulty et al., 2008; Rothwell et al., 1986; Stein et al., 2007). Stein et al. (2007) even went as far as to suggest that RDD may actually be “an epiphenomenon of the resting state.”

However, in these previous studies that showed minimal RDD with ongoing muscle activity, the protocol methods differed greatly from those used in our standard HROC protocol (e.g., position of participant, delivery methodology of stimulation, degree of voluntary activity), leaving us unsure of the true comparability of results. The present study not only confirmed the findings from previous studies but demonstrated that RDD did not occur (regardless of rate, run, sequence, or trial) when the person was providing the low level of soleus background muscle activity generated by standing. This study shows for the first time that it should be possible to perform our HROC protocols at rates as fast as 2 Hz without reducing or destabilizing HR size. A higher rate could increase the number of HR conditioning trials in a session, shorten the session, and/or reduce the required number of sessions. It could thereby accelerate the conditioning process, make the process less demanding for patients, reduce its expense, and increase its practicality. These improvements would increase the number of people, and broaden the breadth of patient populations that could benefit from this important new noninvasive therapy.

### Advantages and Therapeutic Applications

In our current HROC protocols, the recruitment curve and the training trials at our standard rate (0.2 Hz) can take ∼45-60 min to complete, with the patient standing for at least 25 min of that time. This is a challenging task for many, but can be particularly challenging for those with impaired mobility or balance or reduced cognitive endurance/tolerance. A faster rate of stimulus delivery would potentially: (1) allow shorter sessions with much less standing; (2) provide 5-10 times as many conditioning trials (and thus 5-10 times as much feedback to guide HR change) in the same amount of session time; (3) reduce the number of sessions needed for HROC. If a typical session could be reduced from 60 minutes to <25 min, the participant would be less fatigued and better able to then perform skill-specific practice (e.g., upright walking or balance retraining), which drives widespread plasticity. Furthermore, if the dose of conditioning trials is a critical factor for enabling beneficial change in the CNS, the 5-10 times increase in dose provided by faster stimulation might produce a substantially more robust therapeutic effect.

HROC is a powerful and beneficial intervention that could greatly enhance neurologic rehabilitation. However, in today’s healthcare environment, time and billable minutes are major factors in determining the success – and even the survivability – of a neurorehabilitation facility. In translating HROC into the clinical realm, a shorter HROC protocol with equal or greater benefits could add billable skilled-practice time, facilitate insurance reimbursement, and encourage widespread implementation.

### Methodological Issues

As described above, the clinical benefits of increasing the HROC stimulation rate could well be profound. At the same time, these benefits need to be considered in light of some methodological issues that arose during this study.

### M-wave Control

For our HROC protocol, we provide M-wave and HR stability by adjusting stimulus strength as needed to maintain a stable target M-wave size throughout the sessions. This helps to ensure that the same population of primary afferent fibers is excited each time, and thus that the effective strength of the stimulus is stable. The investigator maintains the M-wave target with small changes in stimulation current as needed. With the faster rates (both 1 and 2 Hz), we found that a small adjustment in current predicted from the prior RC was needed to maintain the M-wave target immediately after the participant had a sitting break. To better maintain the target within its range (±20%), initial predicted target currents were recalculated prior to starting the trials and subsequently readjusted as needed. Additionally, at the faster rates controlling the M-wave target through manual adjustment of the stimulation current was more challenging than performing these same adjustments at 0.2 Hz, resulting in some trials being slightly out of the acceptable M-target range (±20%). We are now developing an automatic controller that may provide better control than human operators. However, the delay inherent in feedback-based controllers may still allow occasional trials with out-of-target M-waves.

### Participant Tolerance

Participants reported that the 2-Hz trials were the least comfortable of the three rates assessed. Some participants reported sensory discomfort with the higher rates and slight balance disturbances when not using the minimal upper extremity support of a walker.

### Differences in HR and M-wave due to Stimulation Rate

Our analysis noted a small (∼10%) decrease in the overall HR values for the faster stimulation rates vs. the traditional 0.2-Hz rate. This reduction was within an acceptable range for normal HR variability. It should not affect the ability to effectively perform HROC for therapy. For the HROC therapeutic protocol would always use the same rate; thus, there would be no need to compare HR sizes across different rates.

In addition to the small decrease in the HR values at different rates, there was also an increase in M-wave values at the 1 Hz rate vs the 0.2 Hz. Again, this change was small (increase of ∼14%), falling within our protocol’s ±20% range for acceptable M-wave variation. It should not affect HROC therapy.

Although the small changes in the HR and M-wave values with change in stimulation rate should not impair the ability to perform effective HROC, it does indicate that one should avoid making direct comparisons in HR or M-wave size between the different stimulation rates. For example, pre-testing measurements performed at 1 Hz would not be comparable to post-testing measurements performed at 0.2 Hz.

### Potential effect of interactions between RDD and Pathological State

RDD can reduce or even disappear in certain chronic pathological conditions (e.g., spasticity, NPP, ALS, diabetes) (Aymard et al., 2000; Hultborn and Nielsen, 1995; Ishikawa et al., 1966; Kababie-Ameo et al., 2023; Lamy et al., 2005; Nielsen et al., 1995; Pierrot-Deseilligny and Burke, 2005; Salinas et al., 2022; Schindler-Ivens and Shields, 2004; Worthington et al., 2021; Zhou, X, et al., 2022; Gray et al. 2008) Those with SCI exhibit RDD acutely; however this decreases or disappears when they enter a more chronic phase (Shields et al., 2000). Furthermore, rehabilitation interventions aimed toward restoring function and control can reverse RDD after a stroke and in multiple sclerosis (Kawaishi et al., 2017; León et al., 2023). The disease process, chronicity, and the effects of recovery mechanisms should be considered in the design of an therapeutic intervention (particularly a lengthy intervention) that involves HR measurements at faster rates and/or without BG activity in those with neurological disorders.

### Future Directions

The reliability, rapidity, and magnitude of the HR change produced by an HROC protocol that uses a higher stimulation rate needs testing. We are now undertaking this testing in healthy people and in those with specific chronic neuromuscular disorders. In these studies, we are using the 1-Hz rate which, unlike the 2-Hz rate, provides stable background EMG throughout, and is more comfortable for the participant.

## Conclusion

The present results show that the H-reflex conditioning protocol could potentially be used at stimulation rates as high as 2 Hz. However, 2-Hz stimulation impairs background EMG stability in some participants and is uncomfortable for some. Thus, it may not be optimal for HROC therapeutic protocols. In contrast, the 1-Hz rate is associated with stable background EMG activity and is more comfortable. A 1-Hz HROC protocol could increase times 5 the conditioning trials per session, reduce session duration, and/or reduce the number of sessions. Thus, it might accelerate the conditioning process, make the process less demanding for participants, and facilitate the translation of HROC therapy into widespread clinical use.

## Abbreviations

ALS: Amyotrophic lateral sclerosis
BG: background EMG
CNS: central nervous system
EPOCS: evoked potential operant conditioning system
EMG: electromyography
EXP: exponential
H_max_: maximum H-reflex size
HR: Hoffmann reflex (H-reflex)
HROC: Hoffman reflex (H-reflex) operant conditioning
Hz: hertz
ICC: intraclass correlation
IRB: Institutional Review Board
H_max_: maximum H-reflex size
HRT: H-reflex trials
M-wave: direct muscle response
M_max_: maximum M-wave size
Mr: mean rectified
M-target: target M-wave size
NPP: neuropathic pain
RC: recruitment curve
RDD: rate-dependent depression
SOL: soleus
TA: tibialis anterior

## Author Contribution

All authors conceived the study and design together. JAB, DG, and HM performed the experiments. JAB, RLH, JSC, and DEG performed the analysis. JAB, RLH, JSC, and DEG interpreted the results. JAB, RLH, JSC, and DEG drafted the manuscript. All authors edited the manuscript. All authors approved the final version submitted and are accountable for all aspects of the work.

## Acknowledgements

NIH Grant P41 EB018783 (JRW), NYS Spinal Cord Injury Research Board C37714GG (DG) and C38338GG (JRW), VA SPiRE NCT05880251(JAB, DG), Stratton Veterans Affairs Medical Center

## Ethics Statement

We confirm that we have read the Journal’s position on issues involved in ethical publication and affirm that this report is consistent with those guidelines.

