## Supplementary material for "Soleus H-reflex size versus stimulation rate in the presence of background muscle activity: A methodological study": tables and figures

### Supplementary Materials

Demographic information and experimental treatments received for each participant are in Table 1. Note that the run effect was not found to be significant in the statistical models, thus all six combinations of stimulation frequencies were independently tested (e.g., Participant 1 independently participated in sequence ABC and CBA, even though they were conducted on the same day).

Participant Demographics and Protocol Sequence

| Participant | Gender | Age (yr) | Height (in) | Sequence |
|---|---|---|---|---|
| 1 | M | 31 | 63.5 | ABC-CBA |
| 2 | F | 29 | 67 | BCA-ACB |
| 3 | M | 69 | 72 | CAB-BAC |
| 4 | F | 45 | 63 | ABC-CBA |
| 5 | M | 37 | 69 | BCA-ACB |
| 6 | M | 82 | 72 | CAB-BAC |
| 7 | M | 51 | 70 | ABC-CBA |
| 8 | F | 69 | 66 | BCA-ACB |
| 9 | M | 40 | 69.5 | CAB-BAC |
| 10 | F | 82 | 60 | ABC-CBA |
| 11 | M | 69 | 67 | BCA-ACB |
| 12 | F | 24 | 61 | BCA-ACB |
| 13 | F | 26 | 63 | ABC-CBA |
| 14 | F | 58 | 68 | BCA-ACB |
| 15 | M | 55 | 73 | CAB-BAC |

Table S1. Demographics and stimulation rate sequence (SFS) for each participant. Stimulation frequencies were counterbalanced in a pseudo-randomized block of three for the run 2 that was reversed for run 2: Condition A = 0.2 Hz; Condition B = 1.0 Hz, and Condition C = 2.0 Hz.

### Supporting information for RDD
### (First 20 trials)

Figure S1 shows the raw HR data with a loess fit (i.e., a nonlinear smoother) to estimate any nonlinear mean trends. No consistent nonlinear trend across all subjects appears to exist. Participant 4 appears to have a nonlinear decrease in HR over the first few trials, but this trend is present across all rates.

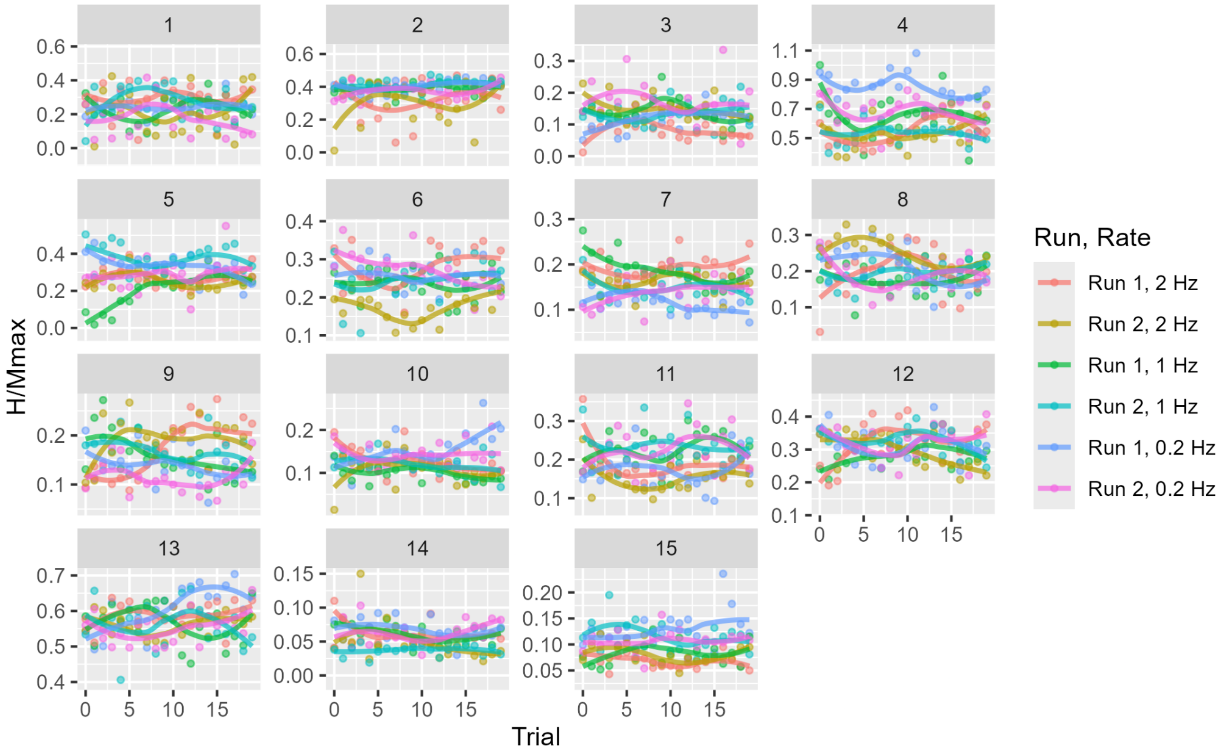


Figure S1: Plot of HR size (normalized to $M_{max}$) vs trial number for each participant by run and rate over the first 20 trials. The mean trend curve is a local least squares smoother ("loess" curve). Note that the y axis scale differs between panels to allow visualization of variability within each subject.

#### Mixed-effects model details for testing presence of RDD

We use mixed-effects models to investigate the presence of non linear trends in HR over the first 20 trials. To capture nonlinearity we modeled log(HR) as the response. We nested participant-level random slopes and intercepts within each treatment condition (i.e., run and rate combination). We fit the following nested (i.e., nested fixed effects) set of models:

- (Full Model, FM) Model with the following fixed effects: main effects of run, rate and trial, all two way interactions between run/rate/trial and a three way interaction between run, rate and trial.
- (Reduced Model 1, RM1) Remove the three way interaction from the FM.
- (RM2) Remove the all two way interactions from Reduced Model 1, keeping only main effects for trial, rate and run.
- (Null model, NM) Model with no fixed effects and only random effects.

The p values for the likelihood ratio test comparisons are as follows, following the nested order: 1) FM vs RM1 (i.e., test the three-way interaction), p=0.72, 2) RM1 vs RM2 (i.e., test the two-way interaction), p=0.64, 3) RM2 vs NM (i.e., test all main effects), p=0.31. In summary, none of the fixed effects are statistically significant.

### Participant-level plots of raw HR, M-wave and BG data (75 Trials)

In Figures S2-S4 are the plots of the raw data for HR, M-wave and BG for all 75 trials. In each figure we observe linear trends within some participants (e.g., in Figure S2, Participant 2 appears to have slight increases in HR across Run and Rate), but no consistent trends across subjects. Similarly, there could be modest differences between run and/or rate within a participant (e.g., in Figure S2, HR appears larger in Run 1, 0.2 Hz compared to other treatments for Participant 4), however that difference is inconsistent across subjects.

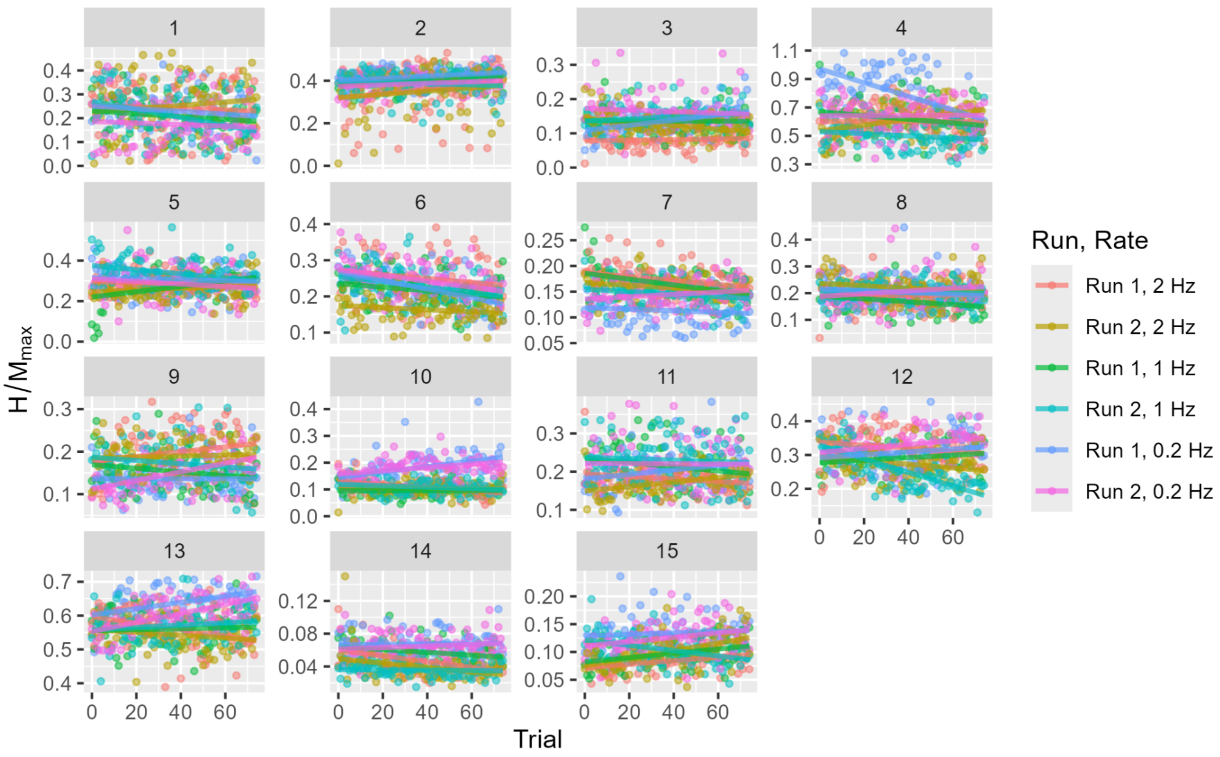


Figure S2: Plot of the raw data for HR size vs trial number for each participant by run and rate. Note that the y axis scale differs between panels to allow visualization of variability within each participant.

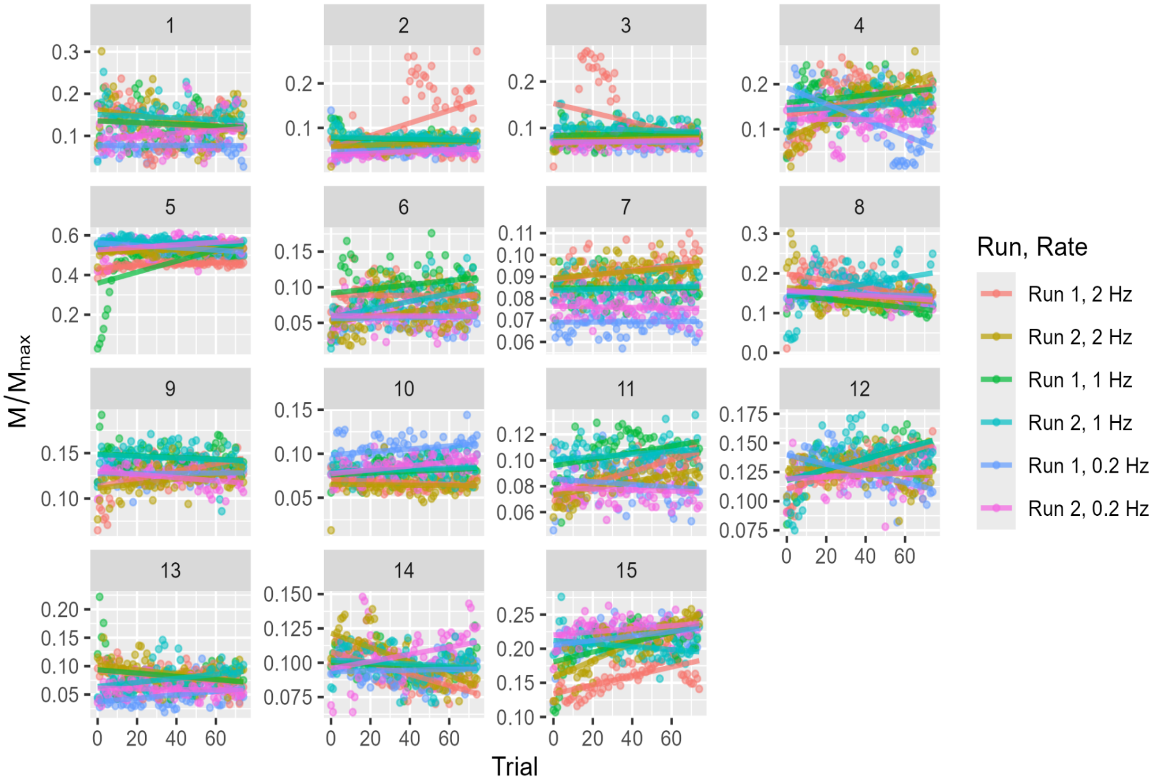


Figure S3: Plot of the raw data for M-wave size vs trial number for each participant by run and rate. Note that The y axis scale differs between panels to allow visualization of variability within each participant.

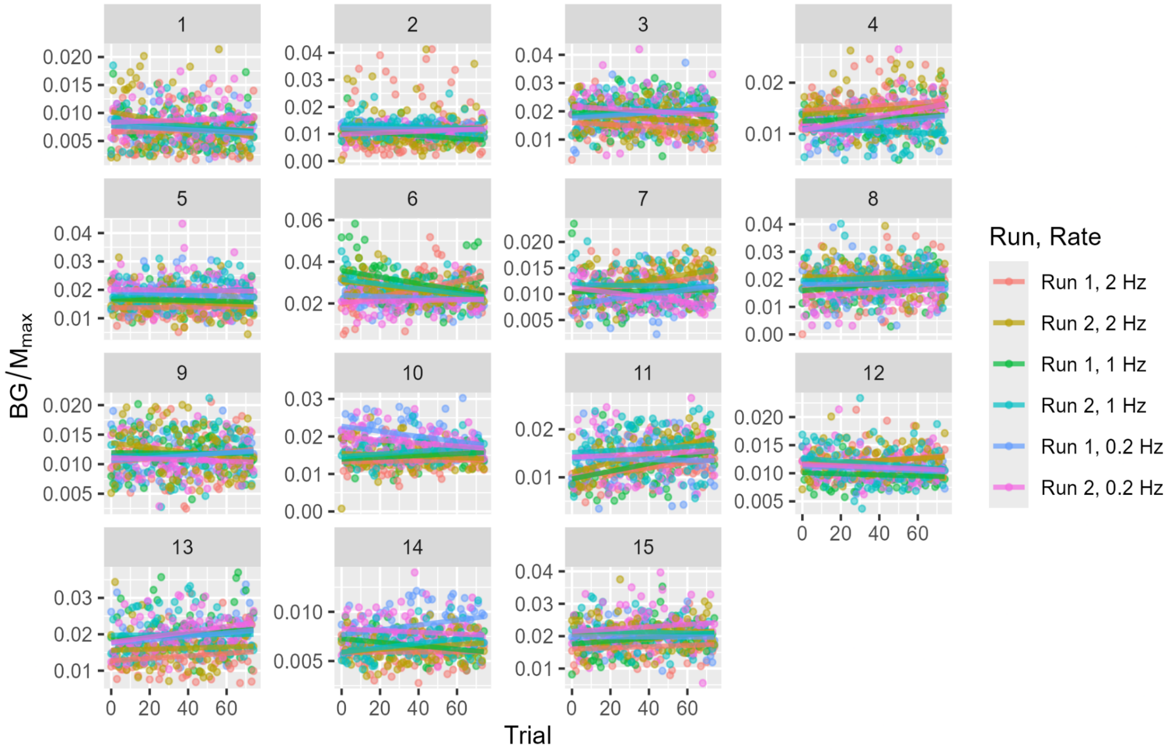


Figure S4: Plot of the raw data of BG vs trial number for each participant by run and rate. Note that the y axis scale differs between panels to allow visualization of variability within each participant.

#### Mixed-effects model details

Analogous to the mixed-effects models employed for detecting RDD during the first 20 trials, we fit mixed-effects models across all 75 trials. We include a linear trend over trial to estimate the overall decrease in HR over the 75 trials. We fit three sets of nested mixed-effects models, with HR, M-wave and BG each normalized to $M_{max}$ as the response variables. Individual models included a random intercept term and random slope for stimulation repetition fit to all combinations of run/rate/participant (Figures S2-S4 show estimates of each random trend line); the fixed effects are outlined below. We compare nested pairs of models using likelihood ratio tests to determine the statistical significance of the fixed effects as follows:

- (Full Model, FM) Model with the following fixed effects: main effects of run, rate and trial, all two way interactions between run/rate/trial and a three way interaction between run, rate and trial.
- (Reduced Model 1, RM1) Remove the three way interaction from the FM.
- (RM2) Remove the all two way interactions from Reduced Model 1, keeping only main effects for trial, rate and run.

- (RM3) Remove trial main effect from RM 2.
- (RM4) Remove run main effect from RM 3, leaving only the rate main effect.
- (Null model, NM) Model with no fixed effects and only random effects.

Table S2 contains the p-values from likelihood ratio tests making the indicated model comparisons (e.g, FM vs RM1 in the first column compares the full model to reduced model 1). A large p-value suggests there is no statistical difference between the two models, thus the terms in the more complex model may be removed with no impact in inference or prediction.

For BG there is no statistical difference between the models (p>0.05 across last row in Table S1), thus the null model - which posits that BG is constant across all trials/runs/rates - explains all the variance in BG after accounting for any random effects. This supports the BG is essentially constant across trial/run/rate.

For HR and M-wave there is evidence of a significant effect or rate, since the comparison of model RM4 (i.e., the model with only rate as a fixed effect) is significantly different from model NM (i.e., the model with no fixed effects), p=0.002 and p=0.02, respectively, for those tests.

| Variable | P value for indicated comparison | | | | |
|---|---|---|---|---|---|
| | FM vs RM1 | RM1 vs RM2 | RM2 vs RM3 | RM3 vs RM4 | RM4 vs NM |
| HR | 0.11 | 0.57 | 0.26 | 0.40 | **0.002** |
| M-wave | 0.35 | 0.66 | 0.13 | 0.36 | **0.02** |
| BG | 0.13 | 0.49 | 0.23 | 0.06 | 0.31 |

Table S2: P values from likelihood ratio tests for testing nested pairs of models.

For HR, we performed a post hoc with Tukey correction to determine the significant differences between rates. There are significant differences between 1 Hz and 0.2 Hz (p=0.002) and 2 Hz and 0.2 Hz (p=0.006). The relative increases in Hmr at 0.2 Hz (mean value 0.259) compared to 1 Hz (mean value 0.231) and 2 Hz (mean value 0.233) are 10.0% and 10.8% respectively.

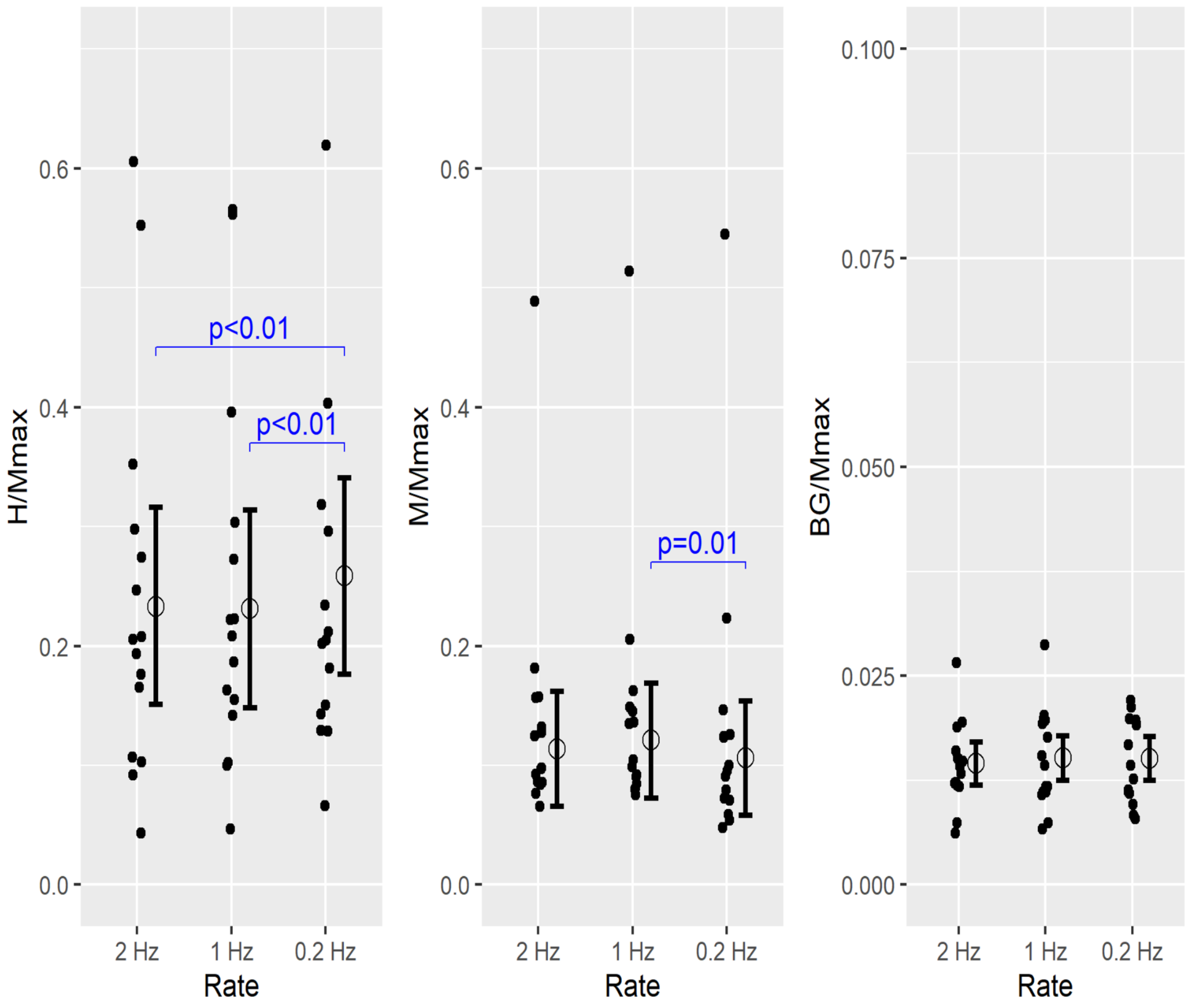


Figure S5. Plots of the mean H-reflex, M-wave, and background EMG (BG) for each participant at different stimulation rates (data averaged across runs and trials). On further exploration of the the averaged data, the mixed-effects models showed that mean HR/$M_{max}$ was significantly smaller at 1 Hz and 2 Hz than at 0.2 Hz (-10.0% and -10.8%, respectively) and that mean M-wave/$M_{max}$ was significantly larger at 1 Hz than at 0.2 Hz (+14%). Although there was no RDD found, there was variability in the mean values with rate, exhibiting the need to always use similar rates and conditions when making comparisons within an experimental design. BG was not affected by stimulation rate.
